# Shotgun Immunoproteomic Approach for the Discovery of Linear B Cell Epitopes in Biothreat Agents *Francisella tularensis* and *Burkholderia pseudomallei*

**DOI:** 10.1101/2021.06.08.447172

**Authors:** Patrik D’haeseleer, Nicole M. Collette, Victoria Lao, Brent W. Segelke, Steven S. Branda, Magdalena Franco

## Abstract

Peptide-based subunit vaccines are coming to the forefront of current vaccine approaches, with safety and cost-effective production among their top advantages. Peptide vaccine formulations consist of multiple synthetic linear epitopes that together trigger desired immune responses that can result in robust immune memory. The advantages of peptide epitopes are their simple structure, ease of synthesis, and ability to stimulate immune responses by means that do not require complex 3D conformation. Identification of linear epitopes is currently an inefficient process that requires thorough characterization of previously identified full-length protein antigens, or laborious techniques involving genetic manipulation of organisms. In this study, we apply a newly developed generalizable screening method that enables efficient identification of B cell epitopes in the proteomes of pathogenic bacteria. As a test case, we used this method to identify epitopes in the proteome of *Francisella tularensis* (Ft), a Select Agent with a well-characterized immunoproteome. Our screen identified many peptides that map to known antigens, including verified and predicted outer membrane proteins and extracellular proteins, validating the utility of this approach. We then used the method to identify seroreactive peptides in the less characterized immunoproteome of Select Agent *Burkholderia pseudomallei* (Bp). This screen revealed known Bp antigens as well as proteins that have not been previously identified as antigens. The present workflow is easily adaptable to detecting peptide targets relevant to the immune systems of other mammalian species, including humans (depending upon the availability of convalescent sera from patients), and could aid in accelerating the discovery of B cell epitopes and development of vaccines to counter emerging biological threats.

## INTRODUCTION

Development of an effective vaccine against a biothreat agent or emerging pathogen is a costly and cumbersome process that can take years to decades to complete. The identification of antigens that stimulate protective immunity against a pathogen represents a significant bottleneck in the typical vaccine development process. Our study addressed the need to accelerate this process by testing the feasibility of a platform for efficient identification of immunoreactive peptides that could be utilized as candidates for development of peptide-based vaccines.

Peptide-based vaccines represent the next generation of vaccines, with great potential to provide rapid protection against biothreats and emerging pathogens. Peptide vaccine formulations consist of multiple synthetic linear epitopes that together trigger immune responses resulting in robust immune memory. This multi-epitope approach can be broadly protective across divergent strains (*e*.*g*., the first universal influenza vaccine to enter phase III clinical trials was a peptide vaccine) and effective for pathogens with complex life cycles (*e*.*g*., several malaria peptide vaccines are currently in clinical trials) (1–3). Due to their lack of complex secondary and tertiary structure, peptides can be easily synthesized, multiplexed into vaccine formulations, and efficiently screened for efficacy. Consequently, peptide-based vaccines represent promising candidates for rapid response medical countermeasures against infectious disease.

Current strategies for epitope identification depend upon detection of epitopes within an individual full-length protein, a low-throughput approach that requires prior knowledge of the antigenic protein and its sequence. Technologies to screen for epitopes at the whole proteome level have been developed (e.g., proteomic microarrays, phage and yeast display); however, these technologies require extensive use of synthetic biology and other time-consuming methodologies (e.g., library construction, peptide/protein array preparation, heterologous protein expression) (3–8). Another major disadvantage of display technologies and use of non-native expression systems is that these methods do not reliably replicate the native properties of the antigenic proteins, including their post-translational modifications, which can lead to inaccurate results.

In this study, proteome-wide screening for linear B cell epitopes was achieved using native proteomes isolated from the pathogen of interest and convalescent sera from infected animals. This strategy holds several advantages over the currently available methods for epitope discovery: It does not require prior knowledge of antigenicity or antigen structure, and obviates need for complex and laborious experimental techniques such as preparation of display libraries and heterologous protein expression. Our approach was designed to enable identification of the protein antigen and, importantly, the antigenic regions within the identified antigen, such that these short linear peptides can be immediately synthesized and tested for efficacy in vaccine formulations.

In this study, we focused on two intracellular bacterial pathogens, *Francisella tularensis* (Ft) and *Burkholderia pseudomallei* (Bp), organisms which pose a high risk for misuse as bioweapons and therefore are considered Tier 1 Select Agents by the US Centers for Disease Control and Prevention. The mortality rates of both pathogens are high, and there is currently no licensed vaccine available for either agent (9–11). Humoral immunity plays an important role in developing immune protection to both of these intracellular pathogens, making them good model organisms for the purposes of this study (12–17). In addition, the immunoproteome of Ft has been thoroughly characterized (10,18,19), such that the previously published data could be compared to the datasets generated in our study. We leveraged a merged dataset of 164 previously identified antigens, corresponding to ∼10% of Ft proteome. The Bp immunoproteome is not as well characterized compared to that of Ft: our reference dataset contained only 61 previously identified seroreactive proteins, corresponding to ∼1% of the Bp proteome (20,21). Consequently, the dataset resulting from the Bp screen has revealed many proteins that have not been previously categorized as antigens.

## MATERIALS AND METHODS

### Bacterial strains and culture conditions

*Francisella tularensis* SCHU S4ΔclpB (“Ft-ΔclpB”) was a generous gift from Dr. Wayne Conlan (National Research Council Canada). Stock cultures were prepared by growing Ft-ΔclpB on Chocolate II Agar plates supplemented with hemoglobin and isovitalex (BD 221169) for 48 hours at 37°C. Bacteria were harvested by scraping confluent lawns into Mueller Hinton (MH) broth containing 20% (w/v) sucrose, and stored at -80°C at a concentration 10^8^ - 10^9^ CFU/mL. *Burkholderia pseudomallei* mutant ΔpurM (“Bp82”) was obtained from BEI resources (NR-51280). Frozen stocks were prepared by growing the bacteria to log phase in Luria-Bertani (LB) broth, adding glycerol to achieve 20% (w/v) with the bacteria at a final concentration of 10^8^ - 10^9^ CFU/mL, and storing aliquots at -80°C. For immunizations, the Ft-ΔclpB and Bp82 bacterial stocks were thawed and diluted in sterile phosphate-buffered saline (PBS) to the specified concentrations used for dosing. For protein extraction purposes, Ft-ΔclpB and Bp82 were propagated to log phase in MH and LB broth, respectively.

### Protein extraction and peptide preparation

Ft-ΔclpB and Bp82 were grown to log phase in 300 mL of MH broth or LB broth, respectively, at 37°C with shaking (250 rpm). The bacteria were harvested by centrifugation at 3200 x g for 10 min at 4°C, washed once with 10 mL of PBS, and the pellet flash frozen using dry ice. The bacteria in the pellet were lysed by subjecting them to two freeze-thaw cycles (alternating between room temperature and dry ice). For protein extraction, the lysate was mixed with Bper Complete Bacterial Protein Extraction Reagent (Thermo Fisher Scientific, cat# 89822), and the mixture incubated at room temperature for 15 min with rotational shaking. The mixture was then subjected to two rounds of sonication (1 sec pulses, timed output 10 sec, at 50% power) using a Heat Systems Ultrasonics sonicator (model W-385), and centrifuged at 16,000 x g for 10 min. Proteins were precipitated with acetone and washed twice with ethanol. Air-dried protein pellets were solubilized using 8M urea and Protease Max surfactant (Promega, V2071), then digested with trypsin (Promega, V5111) using the in-solution digestion protocol provided by the manufacturer (Promega, TB373). Completion of the trypsinization reaction was confirmed by gel electrophoresis. The trypsin-digested proteins were filtered using 10K MWCO concentrators (Pierce) at 10,000 x g for 20 min at 20°C, and the filtrates (purified peptides) stored at -20°C.

### Mice and immunizations

Mouse immunization studies were carried out in strict accordance with the recommendations in the Guide for the Care and Use of Laboratory Animals and the National Institutes of Health. Appropriate efforts were made to minimize suffering of animals. All animals were housed in ABSL2 conditions in an AAALAC-accredited facility, and the protocol (Protocol 270, renumbered 284, approved 10/09/2017) was approved by the LLNL Institutional Animal Care and Use Committee (IACUC). For immunization, 6 week-old female specific-pathogen-free BALB/c-Elite and C57BL/6J-Elite mice (Charles River) were injected subcutaneously with 10^6 CFU Ft-ΔclpB (BALB/c and C57BL/6J), or intradermally with 10^7 CFU Bp82 (BALB/c), and boosted at 2 weeks. No adjuvants were used. Matched PBS-dosed controls were included for each injection route. Course of infection was monitored by performing daily health scoring and weight measurements. Mice that developed infection wounds (Ft only) were topically treated with Dakin’s solution to encourage wound healing, and allowed to remain on test so long as they did not meet humane endpoint criteria (any mice with ∼20% body weight loss or overt signs of morbidity were humanely euthanized). Sera from euthanized mice were excluded from analysis due to lack of immunity to the pathogen. Convalescent sera were harvested from resilient mice at 4 weeks post-infection, *via* cardiac puncture terminal bleeding under inhaled isoflurane anesthesia followed by blood fractionation [centrifugation at 3800 x g for 15 min in microtainer serum separator tubes (BD)]. Sera were stored at -80°CC.

### SDS-PAGE and Western analysis

Western analysis was performed to confirm seropositivity of infected mice. Bacterial lysates were prepared using Bper Complete Bacterial Protein Extraction Reagent (Thermo Fisher Scientific, cat# 89822), combined with Laemmli loading buffer (BioRad), and boiled at 95°C for 5 min. Samples were loaded onto 4-15% acrylamide gels (Mini-Protean TGX, BioRad) and separated by electrophoresis at 120 V for 1 hr. The proteins were transferred from the gels to nitrocellulose membranes (BioRad). Membranes were blocked with Tris-buffered saline plus 0.05% Tween 20 (TBS-T) plus 5% nonfat dry milk, at room temperature for 1 hr or at 4°CC for 16 hrs. The membranes were hybridized with mouse sera at 1:500 dilution in TBS-T plus 5% milk, at room temperature for 2 hrs; washed three times with TBS-T; and then incubated with goat anti-mouse antibodies conjugated to HRP (Pierce, prod#1858413), at 1:5000 dilution in TBS-T plus 5% milk, at room temperature for 1 hr. After three TBS-T washes, the membranes were developed using SuperSignal™ West Pico PLUS Chemiluminescent Substrate (Thermo Fisher Scientific).

### Enzyme-linked immunosorbent assay (ELISA)

ELISA was performed to assess the level of seropositivity of infected mice. Wells were coated with bacterial lysates and incubated at 4°C for 16 hrs. After three washes with PBS plus 0.1% Tween-20 (PBS-T), sera from infected mice diluted to 1:100 with PBS were added to the wells and incubated at room temperature for 1 hr. Following four PBS-T washes, the wells were incubated for 1 hr with Recombinant Protein A/G peroxidase (Pierce, cat#32490) diluted at 1:5000 with PBS. After four PBS-T washes, 1-Step ABTS Substrate Solution (cat# 37615) was added, and after 15 min incubation any colorimetric changes in the wells were detected using a microplate reader (Tecan M200 Pro).

### Affinity purification of immunoreactive peptides

Magnetic beads coated with protein G (Invitrogen, cat#10007D) were used to capture antibodies from sera from infected mice, following the manufacturer’s “Dynabeads Protein G immunoprecipitation” protocol (MAN0017348). The antibody-coated beads were then incubated with purified peptides at room temperature for 45 min. Following three PBS washes, immunoreactive peptides were eluted from the beads using citrate buffer (pH 3). Input, unbound, and eluate fractions were flash frozen with dry ice and stored at -20°C. As a negative control, antibodies from uninfected (PBS treated) mice were used to detect any background resulting from nonspecific binding of peptides to beads or antibodies.

### Mass spectrometry (MS)

The input, unbound, and eluate fractions recovered from antibody-coated beads (see preceding section) were desalted using an Empore SD solid phase extraction plate; lyophilized; reconstituted in 0.1% TFA; and analyzed *via* LC-MS/MS by MS Bioworks (Ann Arbor, Michigan), using a Waters M-Class UPLC system interfaced to a ThermoFisher Fusion Lumos mass spectrometer. Peptides were loaded on a trapping column and eluted over a 75 μm analytical column at 350 nL/min. Both columns were packed with Luna C18 resin (Phenomenex). A 2 hr gradient was employed. The mass spectrometer was operated in a data dependent HCD mode, with MS and MS/MS performed in the Orbitrap at 60,000 FWHM resolution and 15,000 FWHM resolution, respectively. The instrument was run with a 3 sec cycle for MS and MS/MS.

### MS data processing

Data were analyzed using Mascot (Matrix Science) with the following parameters: Enzyme: Trypsin/P; Database: UniProt *F. tularensis* SCHU S4 or UniProt *B. pseudomallei* strain 1026b (forward and reverse appended with common contaminants and mouse IgG sequences); Fixed modification: Carbamidomethyl (C); Variable modifications: Oxidation (M), Acetyl (N-term), Pyro-Glu (N-term Q), Deamidation (N/Q); Mass values: Monoisotopic; Peptide Mass Tolerance: 10 ppm; Fragment Mass Tolerance: 0.02 Da; Max Missed Cleavages: 2; Mascot DAT files were parsed into Scaffold Proteome Software for validation, filtering and to create a non-redundant list per sample. Data were filtered using 1% protein and peptide FDR and requiring at least one unique peptide per protein.

### Bioinformatic analysis

Each experiment typically consisted of three sets of data: “Input” (total bacterial peptides without affinity purification), “Control” (peptides purified from beads coated with antibodies from uninfected mice), and “Experiment” (peptides purified from beads coated with antibodies from infected mice).

LC-MS/MS data were analyzed at the peptide level based on the Total Ion Current (TIC, total area under the MS2 curve), rather than rolling up peptide scores into a protein abundance metric as would be done in standard proteomics. Input datasets were first normalized against each other based on median ratios for the peptides occurring in every Input dataset. The more sparse Control and Experiment datasets were then normalized against their respective Input dataset based on median ratios as well. Since each animal can be expected to raise a different set of antibodies, we counted how often peptides occurred more abundantly in the experiment vs control, rather than focusing on the average log fold change in abundance. Each peptide was assigned an enrichment score, by adding +1, 0, or -1 based on whether the experimental peptide level was greater than, equal to, or lower than the control level in each experiment. Statistical significance was evaluated by randomizing this matrix of +1/0/-1 values.

Average Amino Acid Conservation Scores (AAACS) were calculated using the ConSurf web server(22) with default parameter values, using near full-length protein structure homology models from SWISS-MODEL or crystal structures from PDB where available. The AAACS for the peptide is the average conservation score for the residues in the peptides, with negative scores indicating more highly conserved regions (23).

In addition to AAACS, we also scored peptides based on how many complete sequenced genomes of pathogenic *B. pseudomallei* and *F. tularensis* they occurred in, similar to the conservation analysis in EpitoCore (24). We downloaded proteomes for all 110 *B. pseudomallei* strains with complete genome sequences available through NCBI. For *F. tularensis*, 36 strains with complete genomes were available through NCBI, but several of these corresponded to the less-pathogenic *novicida, holartica* and *mediasiatica* subspecies, so we decided to focus exclusively on the 17 available *F. tularensis* subsp. *tularensis* complete genomes. We identified homologs with ≥ 90% sequence identity to the proteins containing our top scoring peptides in Tables 1 and 2, and then scored each peptide based on how often they had a 100% identical hit in each homolog.

**Table 1:**
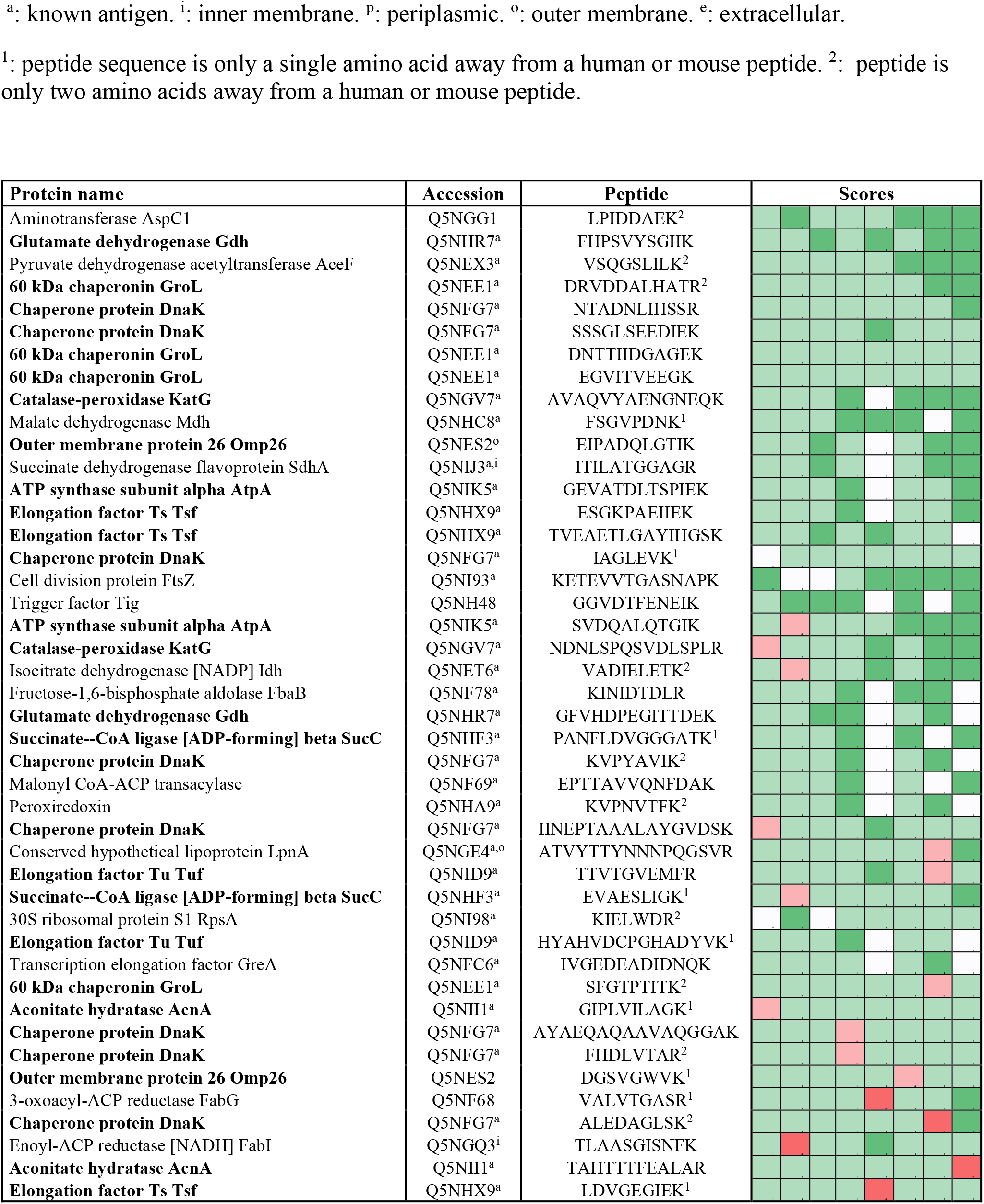
List of top scoring immunoreactive peptides identified for Francisella tularensis. The columns under “scores” indicate whether the peptide was over or underrepresented in each of the 8 experimental samples compared to its control sample. Green: experiment>control. Red: experiment<control. White: peptide undetected in both experiment and control. Dark colors indicate >2-fold difference in relative abundance. Proteins with multiple top scoring peptides are highlighted in bold.

**Table 2:**
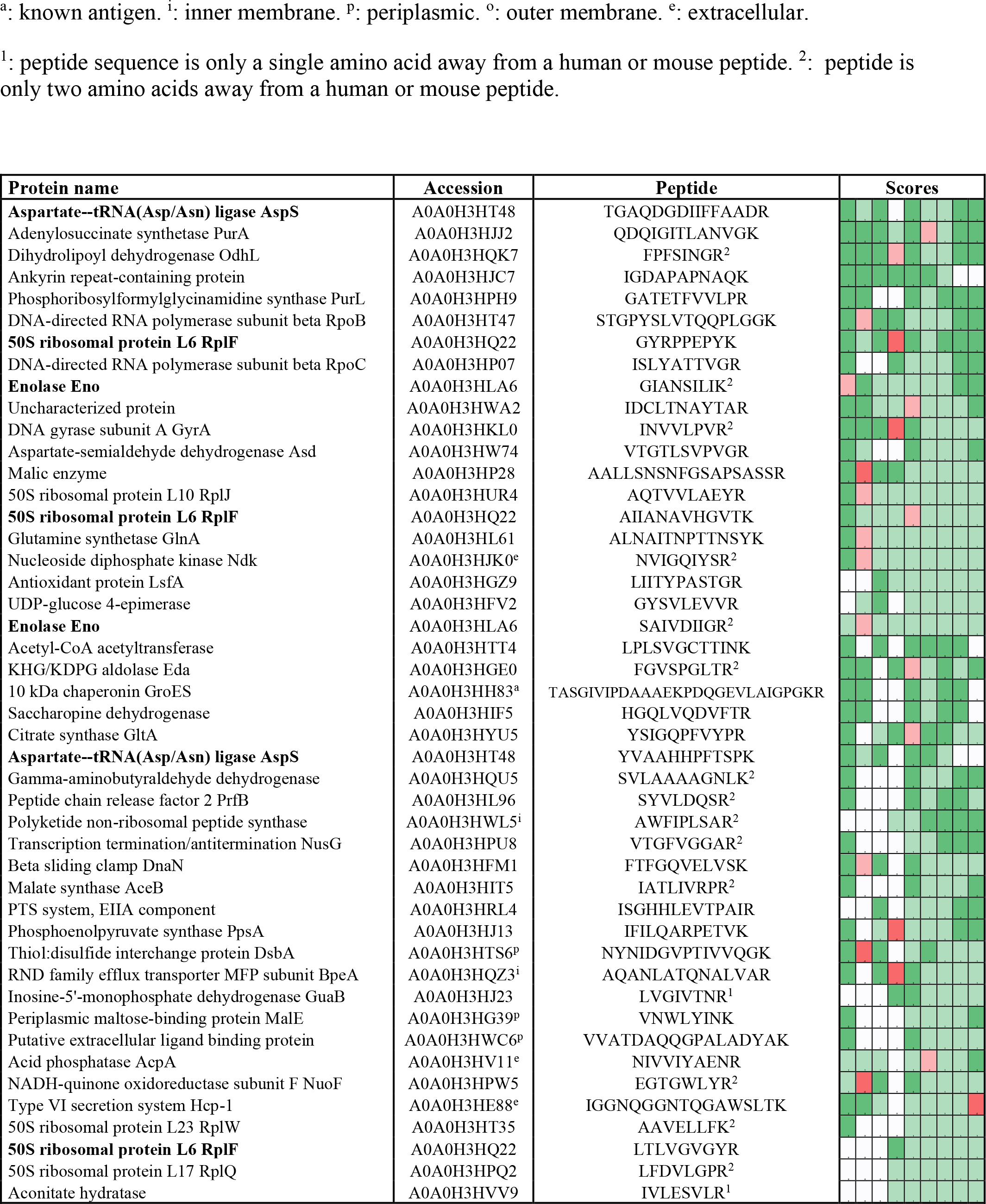
List of top scoring immunoreactive peptides identified for Burkholderia pseudomallei. The columns under “scores” indicate whether the peptide was over or underrepresented in each of the 9 experimental samples compared to its control sample. Green: experiment>control. Red: experiment<control. White: peptide undetected in both experiment and control. Dark colors indicate >2-fold difference in relative abundance. Proteins with multiple top scoring peptides are highlighted in bold.

We used two state-of-the-art computational B cell epitope prediction tools to evaluate all of the peptides in our proteomic data that match the proteins in Tables 1 and 2. Peptides were submitted to the iBCE-EL web server for scoring (25). In addition, proteins were submitted to the Bepipred Linear Epitope Prediction 2.0 tool on the IEDB website (26), and peptides were then scored based on their average predicted residue score. For selected proteins with an available structure model, we also used the Discotope 2.0 web server for prediction of potentially discontinuous B-cell epitopes from protein 3D-structure (27).

## RESULTS

### Overview of immunoproteome screen

In this study, we tested the feasibility of proteome-wide screening for linear B cell epitopes using peptide extracts from target bacteria and sera from infected animals. The method requires: (1) isolation of peptides from lysates generated from the target bacteria; (2) challenge of the host (in this case, mouse) with the target bacteria, followed by collection of convalescent serum; (3) mixing of the bacterial peptides and convalescent serum, to allow peptide antigens to bind to their cognate antibodies in the serum; and (4) recovery of bound peptides for identification through mass spectrometry (Figure 1). We applied this method to two bacterial Select Agent pathogens: *Francisella tularensis* and *Burkholderia pseudomallei*. Infection with attenuated strains of these pathogens [*F. tularensis* SCHU S4ΔclpB and *B. pseudomallei* ΔpurM (strain Bp82)] has been shown to stimulate development of protective immunity against their corresponding fully-virulent parental strains (*F. tularensis* SCHU S4 and *B. pseudomallei* K96245, respectively) (28,29), suggesting that convalescent sera recovered from hosts infected with these attenuated pathogens must contain protective antibodies.

**Figure 1:**
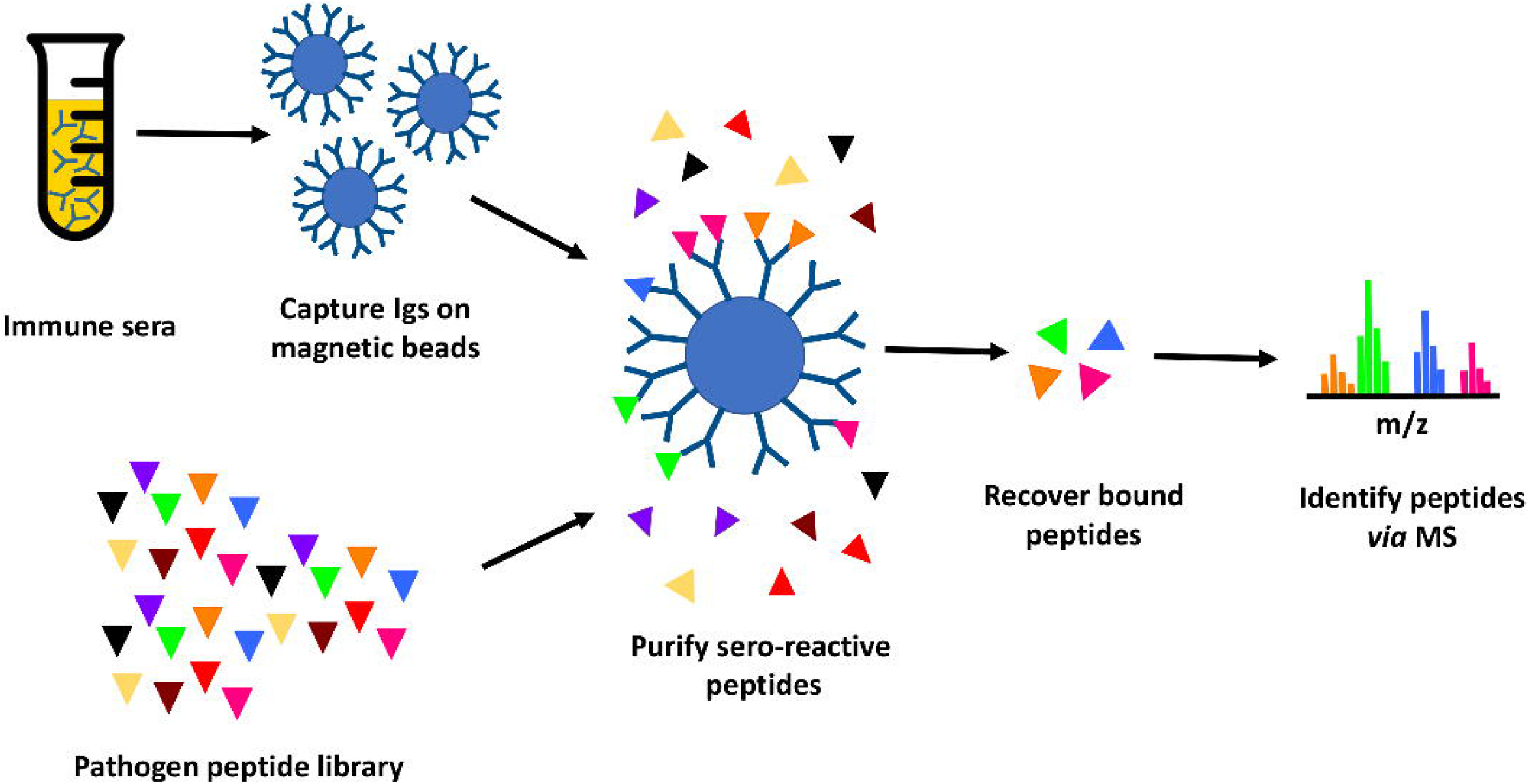
Immunoproteome screening workflow. Schematic overview of high throughput approach for identification of seroreactive peptides in the proteomes of pathogens.

Briefly, proteins purified from pathogen lysates were digested with trypsin to generate a peptide library. Mice were infected with a sublethal dose of Ft-ΔclpB or Bp82, and immune status assessed through measurement of seroreactivity to pathogen lysate *via* enzyme-linked immunosorbent assay (ELISA) or Western blot analysis (Figure 2). Antibodies purified from the convalescent sera of infected mice were immobilized on magnetic beads and then incubated with pathogen-derived peptides to allow formation of antigen-antibody complexes. Peptides recovered from the immobilized antibodies were identified *via* liquid chromatography coupled with tandem mass spectrometry.

**Figure 2:**
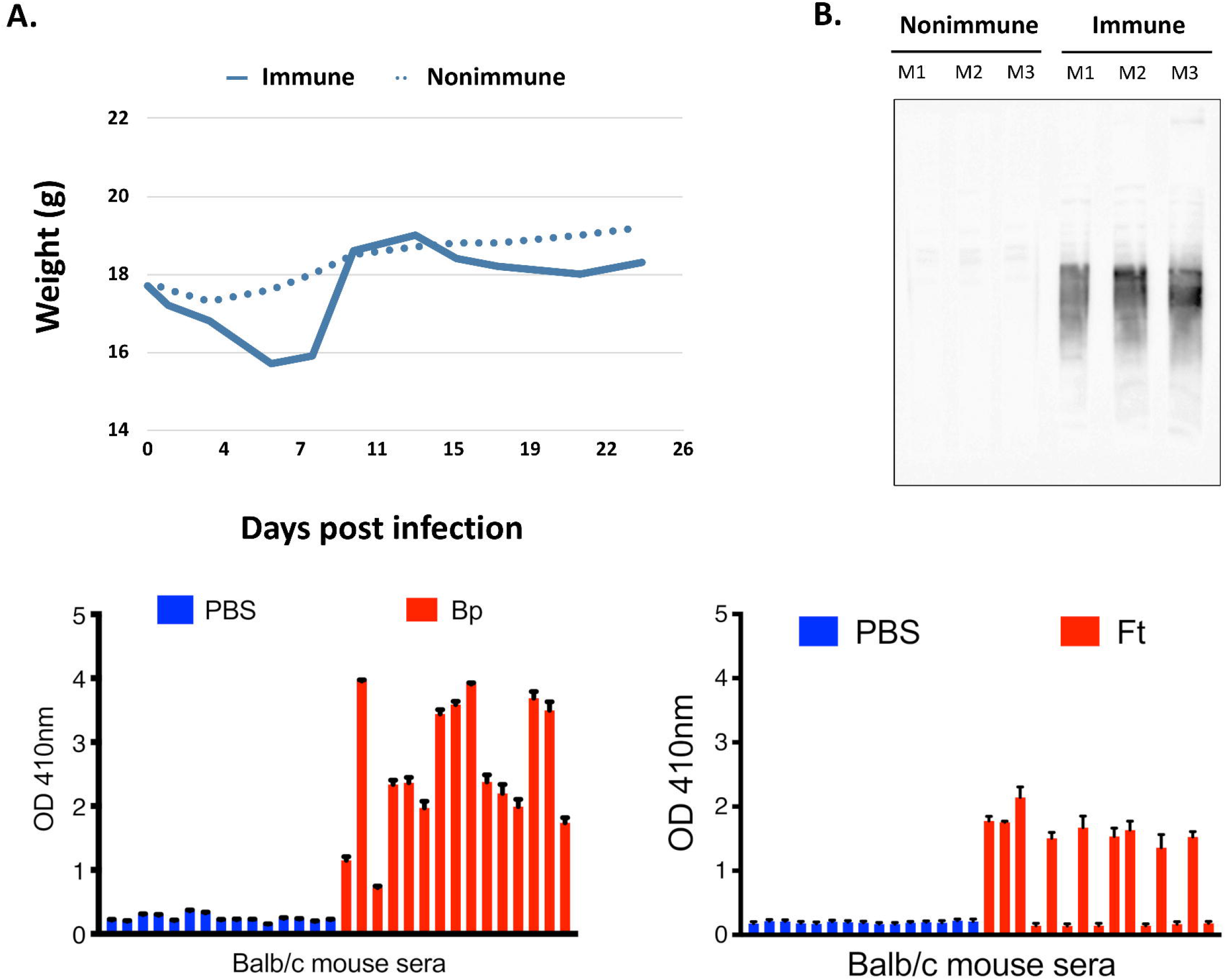
**A**. Representative course of mouse infection to obtain immune sera. Mice were infected with a sublethal dose of Bp and their weight monitored. The degree of weight loss correlates to the amount of antibodies detected in the sera. **B**. Representative Western blot of immune sera vs. non-immune sera. Bp protein lysates were analyzed by Western blotting using sera from infected and uninfected mice (Mouse 1–3). Antibodies from sera with the strongest signal are purified in this study and used to screen for immunogenic peptides. **C**. Representative ELISA results obtained from mice infected with Bp and Ft (red) in comparison with uninfected mice (PBS-treated mice, blue). Sera of some mice infected with Ft did not yield positive results because Ft infection led to lethal outcome and mice had to be euthanized during the course of immunization. Graphs represent two technical replicates for sera collected from each mouse.

### Bioinformatic identification of enriched antigenic peptides

The peptides recovered from infected mice (Experiment peptidome) were compared to those recovered from mock-infected mice (Control peptidome); a total of 8 pairs of peptidomes were collected for Ft, and 9 pairs for Bp. For Ft, we found that 44 of the recovered peptides had an enrichment score of 6 or greater, whereas only 20.1 +/-6.1 peptides would be expected at random (p=5×10^−5^). For Bp, 46 peptides had an enrichment score of 6 or greater, whereas only 17.8 +/-4.3 peptides would be expected at random (p=3×10^−12^). The enriched peptides included those derived from a number of protective antigens identified in previous studies, as well as predicted outer membrane and extracellular proteins (Tables 1 and 2). There were many examples of multiple enriched peptides originating from the same protein (highlighted in bold in the tables), a further indication that enrichment was not random but rather due to immune response to a discrete set of bacterial proteins.

Note that we used C57BL/6J mice for two of the eight Ft experimental samples, because of previously reported differences in protection and antibody response after immunization of C57BL/6J and BALB/c mice with Ft-ΔclpB by Twine *et al* (30). Analyzing the BALB/c Ft samples separately yielded a very similar set of results as in Table 1, but with lower p-value for the enrichment due to the smaller number of samples (results not shown). Therefore, we decided to combine the data and focus on antibody responses in common between both strains of mice. Although Twine *et al* reported an antibody response against chaperonin protein GroL only in BALB/c mice, our data shows that there are several GroL epitopes that are enriched in samples from both mouse strains (see Table 1 and Figure 4).

Immunoproteomics analysis of the antibody response to *F. tularensis* using human or mouse sera has identified 164 antibody targets out of a total of 1667 proteins (∼10% of the entire Ft proteome) (10,18,19). Out of the 1923 peptides that have hits in at least two Ft datasets, 876 peptides match known antigenic proteins. Given those numbers, we would expect only 20 such peptides to show up at random in our list of 44 in table 1, but instead we observe that 38/44 peptides in the list correspond to known antigens - and almost two-fold enrichment (p=2.79×10^−9^). The immune response to *B. pseudomallei* has not been studied in as much depth as for *Francisella*. So even though Bp with 6203 protein coding genes has a genome that is more than three times as large as that of Ft, we found only 61 known antigens identified in previous studies (20,21) (∼1% of the entire proteome). Our list of 47 top Bp peptides in Table 1 includes one known antigen, which does not qualify as a statistically significant enrichment primarily because of the much smaller total number of known antigens for Bp.

Prioritizing highly conserved epitopes is a critical consideration for vaccine development, as highly conserved epitopes can induce broadly protective immunity, and reduce the risk that emergence of pathogen variants will render the vaccine ineffective (31). ∼90% of the top scoring peptides were found to be present in 90% or more of the fully sequenced pathogenic *F. tularensis* and *B. pseudomallei* strains (see Supplementary tables S1 and S2). In addition, we can target peptides that show even deeper evolutionary conservation based on their Average Amino Acid Conservation Score (AAACS), reflecting parts of the protein that may be important for its function (22) (see Supplementary tables S1 and S2).

Note that while some of the proteins in Tables 1 and 2 have homologs in human and mouse (e.g. mitochondrial DnaK), the peptides recovered here are unique to the bacterial versions. Peptides that are only one or two amino acids different from human or mouse versions are likely less suitable as vaccine candidates and are marked with a subscript 1 or 2 respectively in the tables. For vaccine design, we may also want to prioritize peptides which do not tend to occur in healthy human microbiome.

Figure 3 shows the 46 Ft DnaK peptides that were detected in at least two Experiment samples, including the 8 that are in our list of 44 enriched Ft peptides (Table 1).

**Figure 3:**
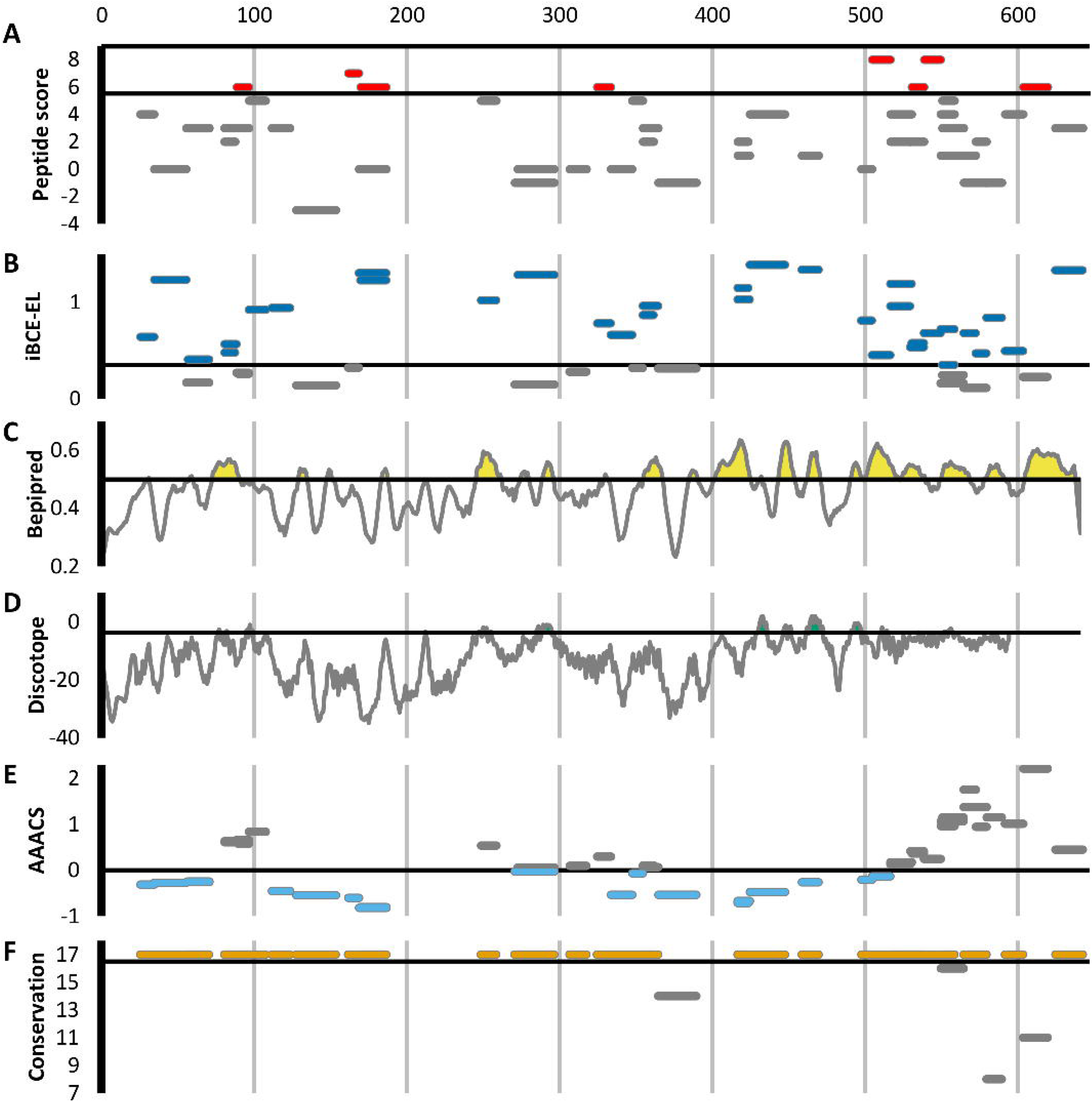
Scoring for the 46 F. tularensis DnaK peptides detected in at least two Experiment samples. Each horizontal line segment indicates the position of a peptide along the length of the 642aa DnaK protein, and its vertical position within each figure panel indicates its score for the metric indicated. The default score threshold suggested for each tool is shown with a horizontal line. **A**. Peptide enrichment score based on our proteomics results. An enrichment score of 8 indicates that the peptide was detected in greater abundance in all 8 Experiment samples relative to their respective Control samples. The threshold for inclusion in Table 1 was an enrichment score of ≥ 6. **B**. B-cell epitope prediction score generated using iBCE-EL. Peptides scoring >0.35 were predicted to be likely B-cell epitopes. **C**. B-cell epitope prediction score generated using Bepipred 2.0. The per-amino acid scores are indicated by the line graph. Regions of the protein scoring >0.5 were predicted to likely contain B-cell epitopes. **D**. B-cell epitope prediction score generated using Discotope 2.0. The per-amino acid scores are indicated by the line graph. Regions of the protein scoring >-0.37 were predicted to likely contain B-cell epitopes. **E**. Average Amino Acid Conservation Score (AAACS) based on Consurf analysis. Lower scores indicate greater degrees of evolutionary conservation. **F**. Number of fully sequenced F. tularensis subsp. tularensis genomes (17 analyzed) in which each peptide occurs.

Lu et al. (32) used hydrogen/deuterium exchange–mass spectrometry (DXMS) to experimentally identify one discontinuous and four linear B-cell epitopes for a selection of mouse monoclonal antibodies against GroL. Figure 4 shows the 32 Ft GroL peptides that were detected in at least two Experiment samples in our study, including the 4 that are in our list of 44 enriched Ft peptides (Table 1). Note that one of these 4 peptides (DNTTIIDGAGEK) overlaps with a linear epitope (NTTIIDGAGEKEAIAKRINVIK) and a discontinuous epitope (SEDLSMKLEETNM— NTTIIDGAGEKEAIA), while a second enriched peptide (EGVITVEEGK) is directly adjacent to another of the linear epitopes (FEDEL). According to the Immune Epitope Database (IEDB) (33), these are the only experimentally validated B-cell epitopes for Ft. IEDB also lists four *B. pseudomallei* antigens that have been assayed for B-cell epitopes, none of which overlap with the proteins in Table 1.

**Figure 4:**
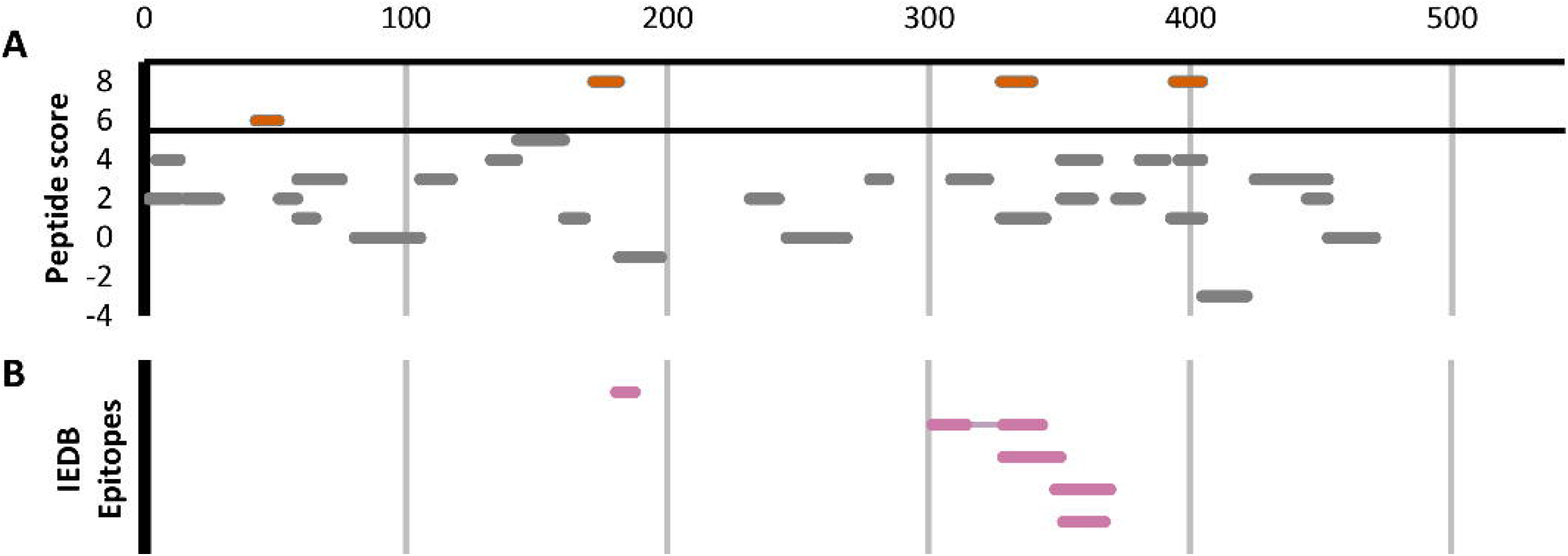
The 32 F. tularensis GroL peptides detected in at least two Experiment samples. Horizontal line segments indicate the position of each peptide along the length of the 544aa GroL protein sequence. **A**. Peptide enrichment score based on our proteomics results, with a score of 8 indicating that the peptide was found in greater abundance in all 8 Experiment samples relative to their respective Control samples. The threshold for inclusion in Table 1 was a score of ≥ 6 or better. **B**. B-cell epitopes identified by DXMS by Lu et al. (32).

## DISCUSSION

We have developed a widely applicable shotgun immunoproteomic method that enables efficient identification of B cell epitopes in the proteomes of pathogens. The results of this study have revealed a significant enrichment of peptides derived from previously identified antigens and vaccine candidates, validating the method’s efficacy. This method was designed to identify linear epitopes efficiently without the need of genetic manipulation or other experimental techniques that can be costly and labor intensive. Attenuated strains made the optimization of this proof-of-concept study more efficient; however, the availability of an attenuated strain for the target organism does not represent a limitation, as our strategy could be applied to fully virulent strains of pathogens as well. Although the present study was performed using a mouse model, the workflow could be easily adapted to detecting targets relevant to the human immune system, using convalescent sera from patients.

Utilizing peptide antigens for vaccine development has several advantages over typical vaccine development efforts. First, peptide vaccines represent a safer alternative to traditional vaccines, because the vaccine formulation is defined and contents are fully synthetic. Second, peptide vaccines have the potential to decrease the cost and production timeline, due to ease of synthesis and recent advances in improved peptide stability (3,34,35). In addition, once antigenic peptides are identified, screening for efficacy could represent a lesser challenge due to the possibility of multiplexing peptides during *in vivo* trials, rather than a one-at-a-time approach.

Among Ft proteins, the present screen identified multiple peptides for two well-characterized antigens, 60kDa chaperonin GroL (Q5NEE1) and chaperone protein DnaK (Q5NFG7). Both chaperonins have been previously implicated in virulence of Francisella (36–38), and are known to induce antibody production in mice and humans (18,39,40). These chaperonin proteins are important for facilitating folding of nascent proteins as well as post-translational modifications. They are also known as heat-shock proteins, as they protect cellular proteins from environmental stresses such as high temperature and low pH (40,41). Although their cellular localization is predicted to be cytoplasmic, they reportedly also associate with membrane proteins and are released into host cells during infection (40,42–44) perhaps contributing to their ability to stimulate various immune functions, including innate immunity, humoral immunity and cell-mediated immunity (36,40,45–48). Heat-shock proteins are good candidates for subunit vaccine design due to their ability to stimulate various immune responses without the need of adjuvant; in fact, both GroL and DnaK have been exploited for vaccine development efforts targeting *Francisella* and other pathogens (32,40,49,50).

Highly virulent Type A *Francisella* strains such as SCHU S4 can bind host plasminogen to the bacterial cell surface where it can be converted to plasmin, a serine protease that degrades opsonizing antibodies, inhibiting antibody-mediated uptake by macrophages (51,52). Among the 25 Ft proteins listed in Table 1, we find at least 3 that are known to be involved in plasminogen binding in *Francisella* or other pathogens, including conserved hypothetical lipoprotein LpnA (Q5NGE4) (52), fructose-1,6-bisphosphate aldolase (Q5NF78) (53), and elongation factor Tu (Q5NID9) (54). These proteins could make for particularly attractive vaccine targets, because if we can interfere with their function before the pathogen has activated its plasmin-mediated antibody evasion, that would make it more susceptible to other antibodies as well.

Among the antigenic peptides identified in the Bp proteome are those belonging to Type VI secretion system component Hcp-,1 and previously identified antigen 10kDa chaperonin GroES (55). Hcp-1 was previously found to be a major virulence determinant in *Burkholderia* and recognized by sera from infected human patients and animals (56–58). Due to this, Hcp-1 has been interrogated as a potential candidate for *Burkholderia* vaccine development (56–58). Additionally, a peptide from an ankyrin repeat-containing protein (A0A0H3HJC) came up as one of the highest scoring peptides in our study. Ankyrin repeats are typically eukaryotic protein domains involved in protein-protein interactions (59), but have been co-opted by many bacterial pathogens as type IV secreted effector proteins to mimic or manipulate various host functions (60).

Recovery of peptides derived from several supposedly cytosolic enzymes may seem puzzling. However several “housekeeping” enzymes are known to be displayed on the surface of pathogens where they play a role in virulence (61). For example, our top scoring peptides from *B. pseudomallei* include two derived from enolase (A0A0H3HLA6). While enolase is primarily thought of as a key glycolytic enzyme, it is also expressed on the surface of a wide variety of bacterial and fungal pathogens, where it interacts with host plasminogen and is associated with invasion and virulence (62). Antibodies against enolase have been detected in a large variety of infectious and autoimmune diseases (63). It is as yet unknown whether enolase plays the same role in Burkholderia, but the protein is predicted to be present both in the cytoplasm and on the cell surface, and its production was found to be upregulated upon exposure to human lung epithelial cells (64). Other housekeeping proteins in our top scoring results whose homologs in other pathogens are known to play a role in adhesion, invasion, or virulence include elongation factor Tu (Q5NID9), malic enzyme/malate dehydrogenase (A0A0H3HP28, Q5NHC8), and fructose-1,6-bisphosphate aldolase (Q5NF78) (61).

Overall, this immunoproteomic workflow has identified numerous peptides mapping to previously identified antigens and subunit vaccine targets, predicted membrane-associated proteins, as well as uncharacterized proteins. The Ft datasets revealed a significant enrichment of peptides belonging to previously identified antigenic proteins in Experiment samples relative to their respective Control samples, providing validation to this approach. Interestingly, several of these known antigens also yielded multiple top scoring peptides in our analysis. Despite the large amount of prior immunoproteomic analysis on Ft, covering ∼10% of the genome, experimentally validated B-cell epitopes are available for only a single protein, and our analysis captures two out of its five known epitopes. Due to the much smaller number of previously identified antigens for Burkholderia, we were not able to tell whether the enrichment in the Bp datasets was significant. More comprehensive immunogenic profiles could be achieved with the use of alternative enzymes with different specificities, since there is a risk of ablating epitopes that contain cut sites recognized by specific enzymes such as trypsin. Alternatively, performing incomplete digestion with one enzyme, or a cocktail of enzymes with different specificities, could improve the yield and diversity of identified epitopes.

Interestingly, we find no significant correlation between the peptides experimentally identified using the method described here, and computationally predicted linear B-cell epitope scores generated by state-of-the-art tools such as Bepipred 2.0 (26) and iBCE-EL (25) (see Supplementary tables S1 and S2), nor any significant correlation between the Bepipred 2.0 and iBCE-EL scores themselves. Accurate computational prediction of B-cell epitopes still poses a major challenge (65), highlighting the value of an unbiased experimental method to screen for antibody targets, as presented here. It is also possible that the tryptic peptides evaluated by this method do not score well as B-cell epitopes by tools such as iBCE-EL, which explicitly take into account sequence features at the beginning and end of the epitope. In cases where the tryptic peptide is too short to be used directly as a vaccine candidate (some are as short as 6 residues), we may be able to use these computational tools to guide us in how to extend the peptide beyond its flanking trypsin cleavage sites.

Further confirmation that the identified sequences are B cell epitopes could be achieved through additional *in vitro* and *in vivo* experimentation (e.g., testing the reactivity of immune sera with synthesized candidate epitopes *via* ELISA or immunization studies). High throughput screening of peptides for efficacy is feasible due to recent advancements in solid phase peptide synthesis (SPPS), which enables efficient and cost-effective production of peptide candidates (3). For immunization studies, pools of multiple peptides could be incorporated into vaccine delivery systems containing adjuvants and T-helper epitopes known to stimulate the induction of adaptive immune response against peptide antigens, as reviewed in Skwarczynski et al (3).

Our immunoproteomic method represents a new tool for precise mapping of linear B cell epitopes. Generation of such immunogenic profiles for pathogens could provide an ample pool of candidates for further experimental validation and efficient vaccine development. Accelerating the discovery of B cell epitopes in the proteomes of pathogens will help fuel the development of peptide-based vaccines that have the potential to provide rapid solutions to biothreat agents and emerging pathogens.

## Data Availability Statement

The mass spectrometry proteomics data have been deposited to the ProteomeXchange Consortium via the PRIDE (66) partner repository with the dataset identifier PXD026300 and 10.6019/PXD026300.

## Author Contributions

PD, NMC and MF contributed to conception and design of the study. NMC performed the in vivo experiments. VL provided laboratory support. VL and MF performed in vitro experimentation. PD performed the bioinformatics analysis. BWS and SSB provided critical input. All authors contributed to manuscript revision, read, and approved the submitted version

## Acknowledgements

We thank Dr. Wayne Conlan (National Research Council Canada) for providing *Francisella tularensis* SCHU S4ΔclpB strain. Bp 82 reagent was obtained through BEI Resources, NIAID, NIH: *Burkholderia pseudomallei*, Strain Bp82 (Δ*purM*), NR-51280. Our thanks go to Michael Ford and MS Bioworks team for help with sample preparation troubleshooting and specialized mass spectrometry analyses. We also thank past and present members of our laboratories - Drs. Sahar El-Etr, José Peña, Amy Rasley, and Emilio Garcia - for useful discussions and critical input.

